# The GINS complex is required for the survival of rapidly proliferating retinal and tectal progenitor cells during zebrafish development

**DOI:** 10.1101/2020.02.09.940767

**Authors:** Máté Varga, Kitti Csályi, István Bertyák, Dóra K. Menyhárd, Richard J. Poole, Kara L. Cerveny, Dorottya Kövesdi, Balázs Barátki, Hannah Rouse, Zsuzsa Vad, Thomas A. Hawkins, Heather L. Stickney, Florencia Cavodeassi, Quenten Schwarz, Rodrigo M. Young, Stephen W. Wilson

**Author notes:** Institute of Ophthalmology, UCL, 11-43 Bath St. London, EC1V 9EL. Corresponding authors: M.V., S.W.W.

## Abstract

Efficient and accurate DNA replication is particularly critical in stem and progenitor cells for successful proliferation and survival. The replisome, an amalgam of protein complexes, is responsible for binding potential origins of replication, unwinding the double helix, and then synthesizing complimentary strands of DNA. According to current models, the initial steps of DNA unwinding and opening are facilitated by the CMG complex, which is composed of a GINS heterotetramer that connects Cdc45 with the mini-chromosome maintenance (Mcm) helicase. In this work, we provide evidence that in the absence of GINS function DNA replication is cell autonomously impaired, and we also show that *gins1* and *gins2* mutants exhibit elevated levels of apoptosis restricted to actively proliferating regions of the central nervous system (CNS). Intriguingly, our results also suggest that the rapid cell cycles during early embryonic development in zebrafish may not require the function of the GINS complex as neither Gins1 nor Gins2 seem to be present during these stages.

## Introduction

In larval zebrafish (*Danio rerio*), extensive cell proliferation during postembryonic stages is observed in proliferative zones of the central nervous system including the ciliary marginal zone (CMZ) at the periphery of the retina and the optic tectum (OT) stem cell niche located along the medial and caudal edges of the tectal hemispheres. Both of these areas contain slow dividing neural stem cells adjacent to fast proliferating transit amplifying (TA) cell populations (Cerveny et al., 2012; Galant et al., 2016; Joly et al., 2016; Recher et al., 2013). The elevated proliferation rate in these cells is sustained by the expression of high levels of genes required for cell cycle progression, including those encoding DNA replication proteins. Transcripts and protein products of genes involved in cell cycle progression, including DNA replication and mitosis, are maternally inherited and stable, supporting the rapid proliferation that occurs during the earliest stages of vertebrate development (Aanes et al., 2011; Alli Shaik et al., 2014; Vesterlund et al., 2011). In developing zebrafish, therefore, a number of recessive mutations in cell cycle linked genes first manifest phenotypes at later stages in neural stem and progenitor cells, causing defects in the proliferative zones of the OT and retina (Amsterdam et al., 2004).

The eukaryotic replication complex (also known as the replisome) is a multi-subunit molecular machine that assembles at replication origins. In this evolutionarily ancient complex, which has well-defined orthologs even in Archea (Xu et al., 2016), the helicase subunit unwinds the DNA strands, while a polymerase and a primase contribute to new strand synthesis. The eukaryotic helicase, the Cdc45-MCM-GINS (CMG) complex, is itself a compound structure comprised of the Mcm2-7 motor subunits, the GINS (Go, Ichi, Nii, San) heterotetramer and Cdc45 (Sun *et al.,* 2015; Yuan *et al*., 2016). As revealed by analyses in budding yeast, the GINS complex tethers the Mcm2-7 subunits and Cdc45 to the replication origin (Takayama *et al.*, 2003). A combination of structural studies demonstrated that the GINS complex forms a horseshoe-shape consisting of a pseudo-symmetrical layered structure of two heterodimers, mediating the interaction of Cdc45 and other components of the replisome with the MCM complex. The GINS complex, together with Cdc45, acts as a scaffold for the N-tier of the helicase and is essential for helicase function (Boskovic *et al*., 2007; Gambus *et al*., 2006; Kamada *et al*., 2007; Sun *et al*., 2015; Yuan *et al*., 2016).

Loss of function of the GINS orthologues *sld5* and *psf1-1* in budding yeast causes growth defects (Takayama *et al*., 2003) and in *Drosophila*, impairment of GINS components causes chromosome condensation and genomic integrity defects (Chmielewski *et al*., 2012; Gouge and Christensen, 2010). Mouse embryos homozygous for mutations in *Sld5* and *Psf1* die around implantation stages (Mohri *et al*., 2013; Ueno *et al.*, 2005) whereas in *Xenopus*, knockdown of *psf2* (the frog ortholog of *GINS2*) results in retinal development defects (Walter *et al*., 2008).

In healthy humans, higher expression of *GINS* orthologs is detected in proliferative tissues relative to differentiated cells (Uhlén *et al*., 2015). Enhanced expression of GINS components is also reported in numerous cancer types, and these genes can serve as important biomarkers for cancer therapy (Kanzaki *et al.*, 2016; Tauchi *et al.*, 2016; Yamane *et al*., 2016). In several cancer cell lines, inhibiting GINS function results in a decrease in proliferation and invasive behaviors, suggesting that the complex may be an important therapeutic target (Liang *et al.*, 2016; Yamane *et al*., 2016; Zhang *et al.*, 2015; Zhang *et al*., 2013).

In zebrafish embryos, expression of CMG components is restricted to proliferative tissues and by 2 dpf a number of these genes are expressed preferentially in the CMZ and OT (Thisse and Thisse, 2004; Thisse *et al*., 2001; Thisse and Thisse, 2005). Although an insertional mutagenesis screen identified mutations in genes encoding several CMG components in zebrafish (Amsterdam et al., 2004), only *mcm5* has been studied in detail. Like many other cell cycle related genes, *mcm5* is expressed primarily in stem and progenitor cells in the retina and OT and mutations in *mcm5* elicit apoptosis of proliferating progenitor cells in those regions (Ryu et al., 2005).

In a recent genetic screen, we isolated several mutants that exhibit apoptosis preferentially in the CMZ and the proliferative regions of the OT at 2 days post fertilization (dpf). Interestingly, after 2 dpf, the apoptotic phenotype recedes and some of these mutant larvae survive for up to 7 dpf. At 5 dpf, however, they all lack the stereotypical laminated architecture characteristic for the OT. Genetic mapping and characterization of one of the mutants linked these phenotypes to a mutation in *gins2*, a gene encoding an essential member of the eukaryotic replication complex.

We show here that the Gins2^L52P^ mutation abolishes CMG function, with highly proliferative cells arresting in the S phase of the cell cycle and as a consequence undergoing apoptosis. Surprisingly *in silico* molecular modeling analyses suggest that the mutation does not completely disrupt the GINS complex completely, but instead induces subtle though significant changes in the interaction surface between Gins2 and Gins4 proteins, which might affect the stability of the complex.

## Results

### Isolation of mutants with excess apoptosis in the retina and the tectum

In an ENU-based genetic screen in zebrafish, we isolated the *u773* mutant line that shows elevated cell death in the eyes and the OT at 2 dpf. Live homozygous mutant embryos show a characteristic dark patch is seen in the OT, and TUNEL staining confirmed that this appearance is due to increased cell death in the tissue (Figure 1E,F). Surprisingly, after 3 dpf there is a reduction in the number of apoptotic cells (Supplementary Figure 1), but while the mutants survive up until 5 dpf, they show smaller eyes and smaller and severely disorganized OT (Figure 1C,D,G-H’). Histological analysis of mutant larvae showed that the eye and OT lack cells within the retinal and tectal progenitor domains (Supplementary Figure 2.)

**Figure 1.**
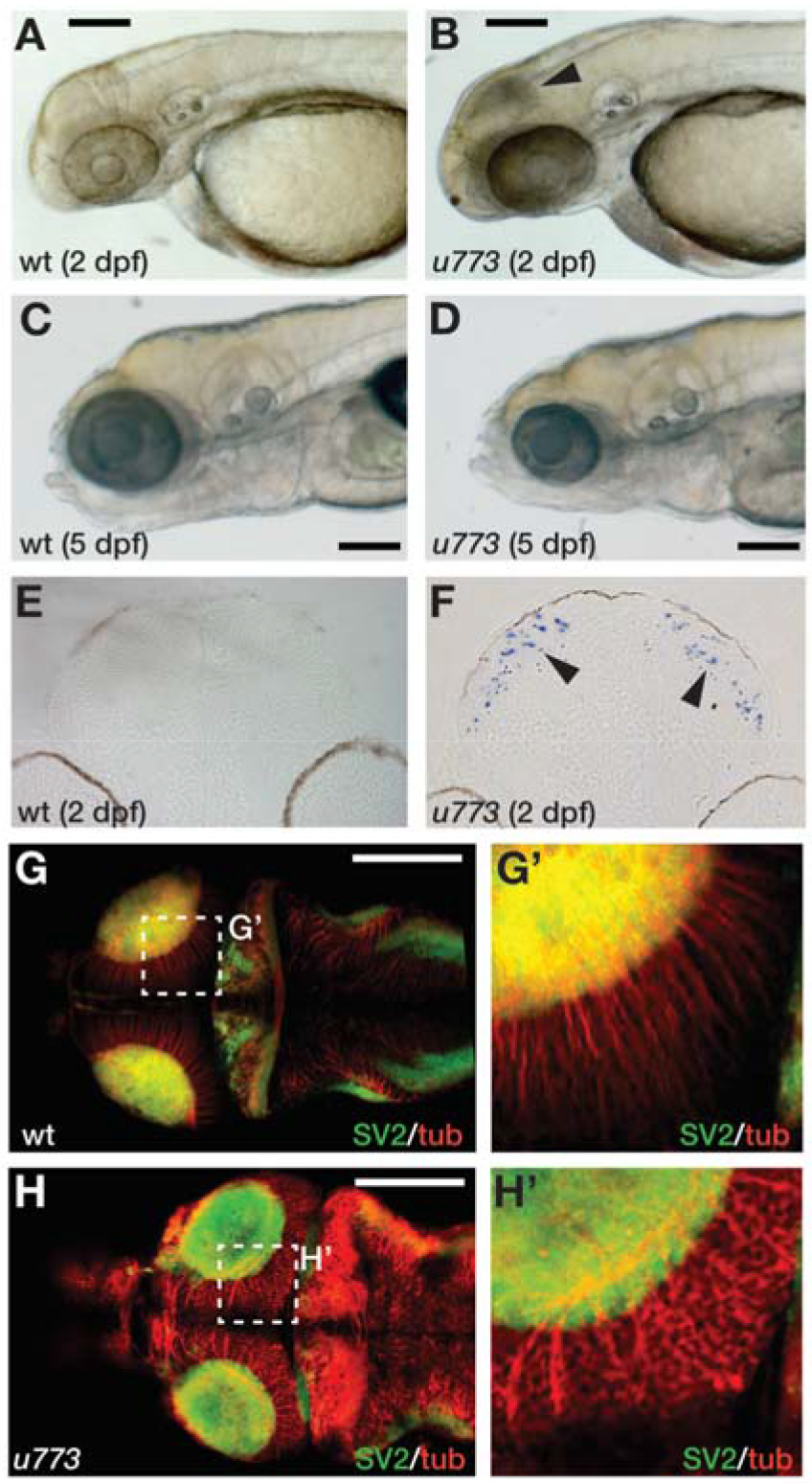
*u773* mutants show tectal apoptosis. A-D) Lateral views of live wildtype (A,C) and *u773* mutant (B,D) embryos at stages indicated. The arrowhead in B indicates dying tectal cells. E,F) Transverse sections of the tecta of *u773* mutants and siblings showing TUNEL labeled apoptotic cells (blue, arrowheads). G-H’) Dorsal views of brains of *u773* mutants and siblings labeled with anti-acetylated tubulin (red) and SV2 antibodies showing organization of cells, processes and neuropil. G’ and H’ show magnified views of the boxes in G and H. Note that in the *u773* mutant, tubulin staining is aberrant. Scale bars: 200 μm.

### The *u773* mutation is in the *gins2* gene

Simple sequence length polymorphism (SSLP) mapping located the *u773* mutation to LG18 between markers z7256 (42.1 cM, MGH panel) and z10008 (44.2 cM, MGH panel) (Figure 2A). Single nucleotide polymorphism homozygosity mapping using the Cloudmap platform (Minevich et al., 2012) on whole-genome sequencing (WGS) data confirmed this chromosomal position (Figure 2B) and showed that one of the genes in the interval, *gins2,* carries a missense mutation (Figure 2C). This c.217T>C mutation in the *gins2* cDNA results in an L52P change in a highly conserved region of the Gins2 protein (Figure 2H).

**Figure 2.**
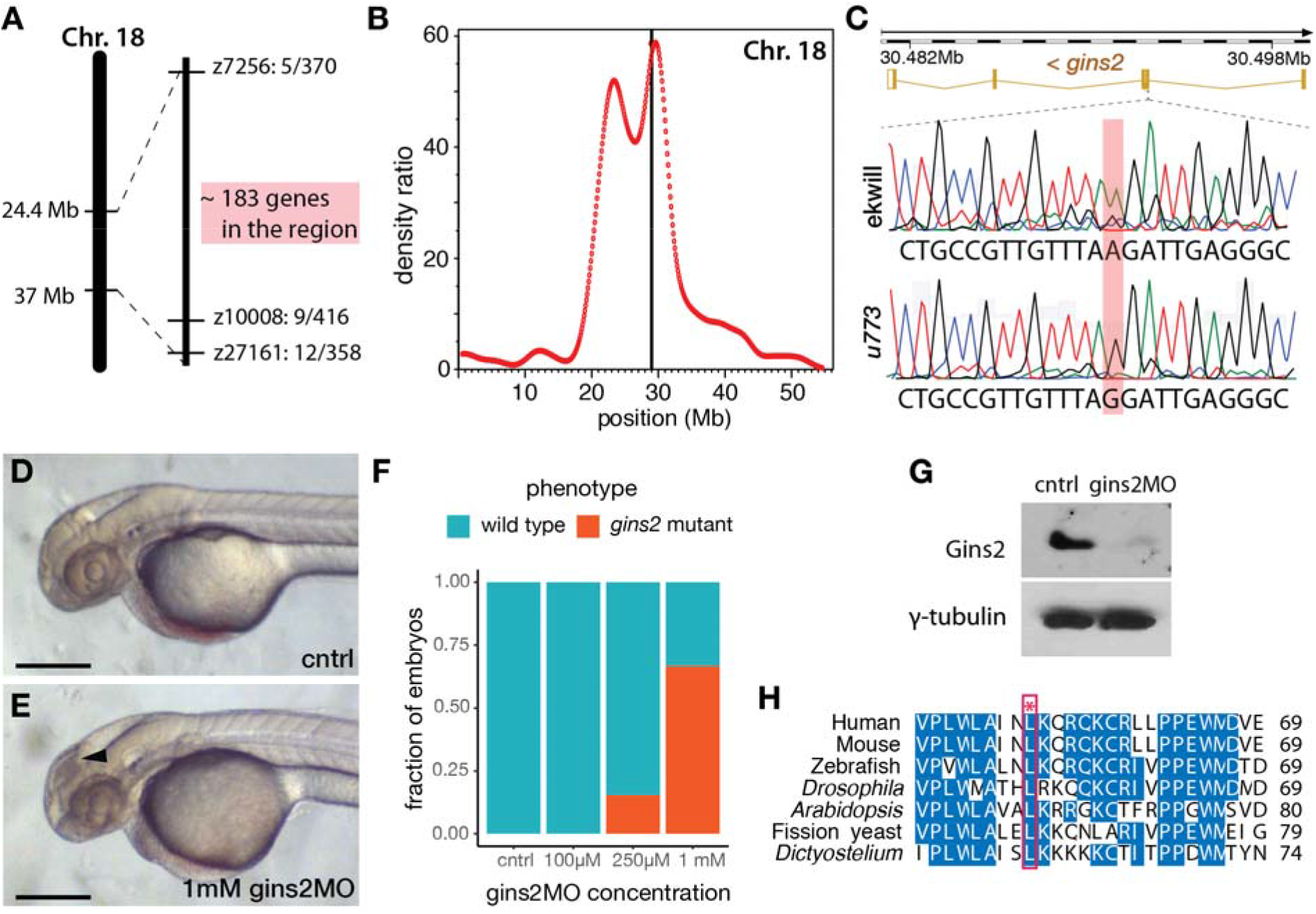
*u773* is a loss-of-function allele of *gins2.* A) Initial region of Chromosome 18 that segregated with the *u773* mutant phenotype. B) WGS mapping plot of SNP homozygosity on Chromosome 18. C) Sanger sequencing results aligned to the reference sequence of *gins2* for *u773* mutants and ekwill controls. (Note that *gins2* is on the reverse strand and the sequence is shown in the reverse direction.) D, E) Phenotype of 2 dpf *gins2MO* injected embryos and uninjected controls (black arrowhead indicates dead cells). Scale bars: 250 μm. F) Graph showing the fraction of morphants showing the *gins2^u773^* phenotype at the indicated *gins2MO* concentrations. G) Protein lysates from uninjected (cntrl) and *gins2*MO-injected 2 dpf embryos were separated with SDS-PAGE and then subjected to Western blotting with Gins2 and γ-tubulin antibodies. H) Multiple sequence alignment of Gins2/Psf2 protein sequences in different species. The red box and asterisk denotes the lysine residue affected in *u773* mutants.

To confirm that the *u773* phenotype results from the mutation in *gins2,* we performed a series of experiments using a synthetic morpholino oligonucleotide that targets the splice-donor site of the first exon of *gins2 (gins2MO).* Injection of the *gins2MO* into wildtype embryos reduced *gins2* mRNA and Gins2 protein levels (Figure 2G, Supplementary Figure 3) and resulted in a concentration dependent cell death phenotype similar to that observed in the OT of *u773* mutants (Figure 2D-F). Therefore, we conclude that the mutation in the *gins2* gene underlies the apoptotic phenotype observed in homozygous *u773* embryos, and that the Gins2^L52P^ mutation likely results in a complete loss-of-function phenotype.

### *gins2* functions cell-autonomously and is expressed from somitogenesis

As a member of the replisome, Gins2 is expected to behave in a cell-autonomous fashion. To address this, we transplanted cells from either wildtype or *gins2*MO-injected embryos into the presumptive eye-field of wildtype embryos. Similar to *gins2^u773^* mutant eyes, *gins2* morphant cells in wildtype embryos exhibited elevated levels of apoptosis (Figure 3A-B’), demonstrating that loss of Gins2 triggers cell death in a cell autonomous manner.

Similarly to genes encoding for the other components of GINS complex (Thisse and Thisse, 2004; Thisse et al., 2001), *gins2* transcripts were not observed before somitogenesis – 12 hpf (Figure 3C-G). At 1 dpf expression was localized to the fast proliferating cell populations throughout the embryo (Figure 3H), and by 2 dpf *gins2* transcription was observed primarily in the CMZ and the proliferative regions of the OT (Figure 3I, I’). This 2 dpf expression pattern is typical to the components of the zebrafish CMG apparatus (Thisse and Thisse, 2004; Thisse et al., 2001).

**Figure 3.**
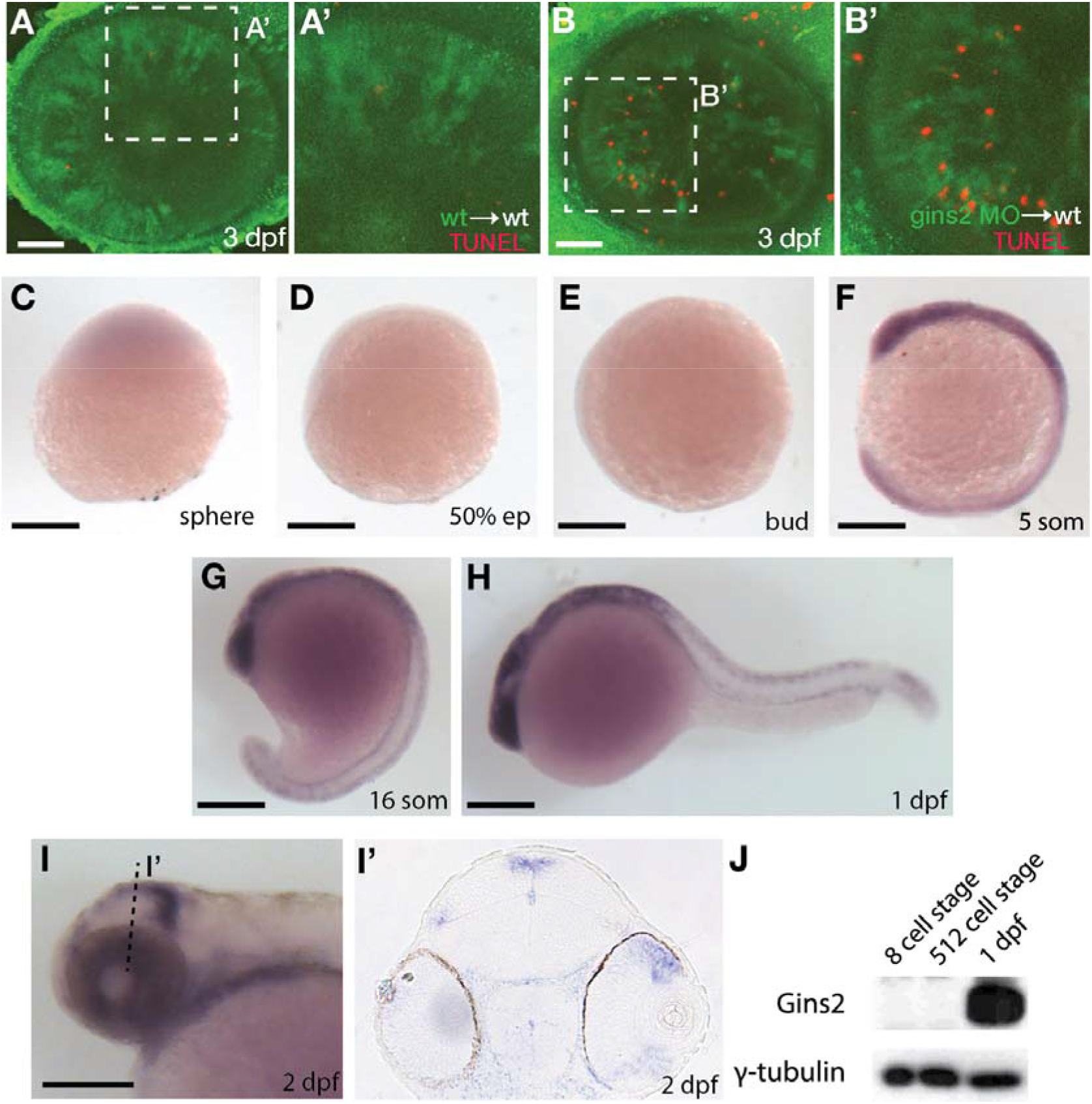
*gins2* functions cell-autonomously and is expressed in proliferating tissues. A,B) TUNEL staining (red) of wildtype host embryos receiving heterologous cell transplants labeled with membrane-tethered GFP (green) from wildtype and *gins2*MO-injected embryos. A’ and B’ show magnified views of the boxes in A and B. C-I’) Lateral views (C-I) and transverse section (I’) of the wildtype embryos assessed for zygotic *gins2* expression with whole mount *in situ* hybridization at the indicated stages. J) Western blot analysis of protein lysates from wildtype embryos probed with Gins2 and γ-tubulin antibodies. Scale bars: (A, B) 50 μm; (C-I) 150 μm.

As Gins2 is considered a canonical component of the eukaryotic replisome, we reasoned that the relatively late expression of *gins2* during development might be a consequence of the presence of a strong maternal component that enables cells to proceed through the early divisions. To our surprise, however, we observed no Gins2 mRNA or protein in early embryos (Figure 3), suggesting that cell divisions before somite stages might occur in a Gins2-independent manner.

### There is no functional redundancy between Gins1 and Gins2

On the basis of observations described above, the reason for the late-onset *gins2* loss-of-function phenotype is not obvious. However, sequence analysis of the distinct proteins that form the GINS complex in Eukaryotes and Archea suggests that they all arose from a single ancestral protein (Kelman and Kelman, 2014; Makarova et al., 2005; Xu et al., 2016). Therefore, one possible explanation for the late-onset, tissue-specific phenotype observed in the *gins2^u773^* mutants could be some functional compensation by other GINS components during zebrafish development. Indeed in budding yeast, overexpression of certain GINS components can rescue thermosensitive mutations in others (Takayama et al., 2003).

To address a possible redundancy between different zebrafish Gins proteins, we generated a *gins1* mutant line using CRISPR/Cas9-based genome editing. The *gins1^elu11^* allele has a 2 bp deletion in its exon 4 (c.250_251delCT), and homozygous *gins^elu11^* mutants show a phenotype comparable to that observed in other mutants of the CMG complex, with characteristic apoptosis in the fast proliferating cell populations of the OT and the retina (Figure 4A,B). Western blot analysis suggests the *gins1^elu11^* allele is not able to produce functional Gins1 protein (Figure 4F), therefore, it is a functional null mutant.

**Figure 4.**
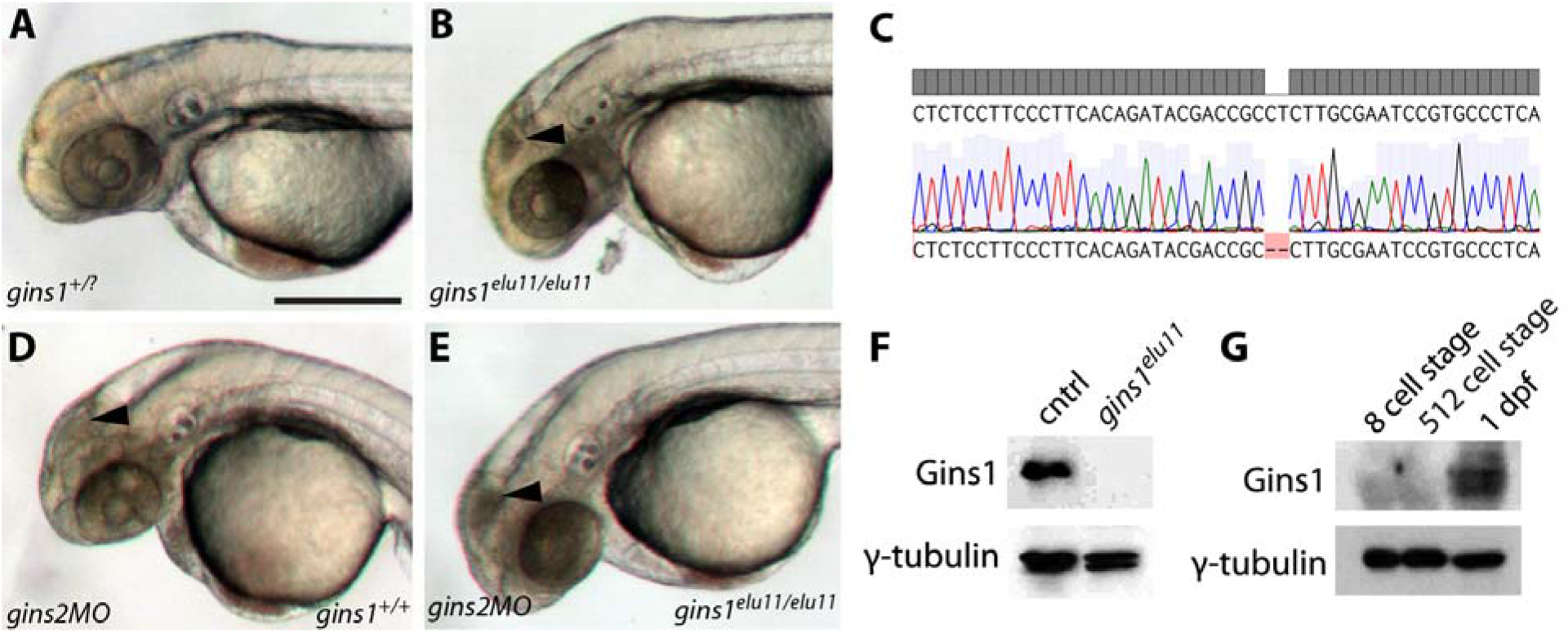
Gins1 deficient and Gins1, Gins2 double-deficient embryos show comparable cell death phenotypes in eyes and tecta. A,B) The *gins1^elu11^* frameshift allele created through genome editing shows retinal and tectal apoptosis at 2 dpf, similar to other CMG mutants. C) Sanger sequencing of *gins1^elu11/elu11^* embryos shows the presence of the homozygous c.250_251delCT mutation. D,E) Phenotype of *gins1* mutant embryos and their siblings injected with 100 μM *gins2*MO. F) Western blot analysis of protein lysates from 2 dpf control and *gins1^elu11/elu11^* embryos probed with Gins1 and γ-tubulin antibodies. G) Western blot analysis of protein lysates from wildtype embryos at the indicated stages probed with Gins1 and γ-tubulin antibodies. Scale bar: 250 μm.

Knockdown of *gins2* in *gins1^elu11/elu11^* homozygous mutant embryos did not reveal enhanced cell death compared to either single mutant: only the aforementioned progenitor populations showed high levels of apoptosis in *gins2MO* injected mutants. Therefore, we conclude that there is little or no functional redundancy between these two GINS components during early zebrafish development, and the late-onset apoptotic phenotype observed in the OT and the retina is likely due to a complete loss of GINS function in single *gins1* or *gins2* mutants.

We also assessed the expression of *gins1* and found that similarly to *gins2,* it starts only during somitogenesis stages, after 12 hpf (Suppl. Fig. 4). Furthermore, maternal Gins1 is also absent during early phases of development, suggesting that the absence of Gins proteins is a general feature of the early zebrafish embryo (Fig. 4G).

### Characterization of cell cycle defects in *gins2* mutants

A mutation in the zebrafish CMG complex *mcm5* gene leads to cell cycle slowdown and arrest, (Ryu et al., 2005), we examined how cells deficient in Gins2 progress through, and exit, the cell cycle. First, we performed BrdU pulse-chase experiments by exposing 2 dpf embryos to BrdU and examining the BrdU content of tectal cells at 3 dpf. In the OT of wildtype embryos, a medial-to-lateral gradient of fluorescence (BrdU content) is observed at this stage (Figure 5A), which reflects the dilution of the BrdU in the nuclei of the proliferating cells during the chase period due to semiconservative DNA replication. In contrast, in the OT of *gins2^u773/u773^* mutants no such dilution was observed (Figure 5B), suggesting that mutant cells in the tectal proliferative zone do not undergo repeated cycles of DNA replication and may stall in S-phase. The apparent increase in BrdU-positive cells is also explained by the failure of the cells in the proliferative zone of the mutant OT to go through rounds of cell cycle following the BrdU pulse.

To understand the nature of cell cycle defects that arise upon the impairment of *gins2* function, we compared the DNA content of cells isolated from the heads of sibling and mutant embryos. Flow cytometric analysis of the DNA content in cells isolated from the head of *gins2* mutants showed a slight reduction in the ratio of 4C cells compared with their siblings (Figure 5C-E). These observations are in concordance with earlier data from untransformed human dermal fibroblasts, where impairment of Gins1/Gins2 reduced the rate of DNA synthesis and delayed the cell cycle (Barkley et al., 2009). As similar experiments performed on retinal cells from *mcm5* mutant zebrafish larvae suggested that cell cycle defects in those mutants could be attributed partly to a block in cell division after DNA replication (Ryu et al., 2005), our results suggest that loss of different CMG components result in slightly different cellular-level phenotypes.

Zebrafish embryos lacking *mcm5* function show jaw developmental defects, with late differentiating cartilages failing to form (Ryu et al., 2005). This is due to the peculiarities of cranial cartilage development, where chondrocytes undergo directional proliferation followed by hypertrophy (Yan et al., 2002). *Gins2^u773^* mutants show similar severely deformed jaw cartilages to those observed in *mcm5* mutants. Quantifications of jaw parameters showed highly significant differences between sibling and mutant tissues (Supplementary Figure 5).

**Figure 5.**
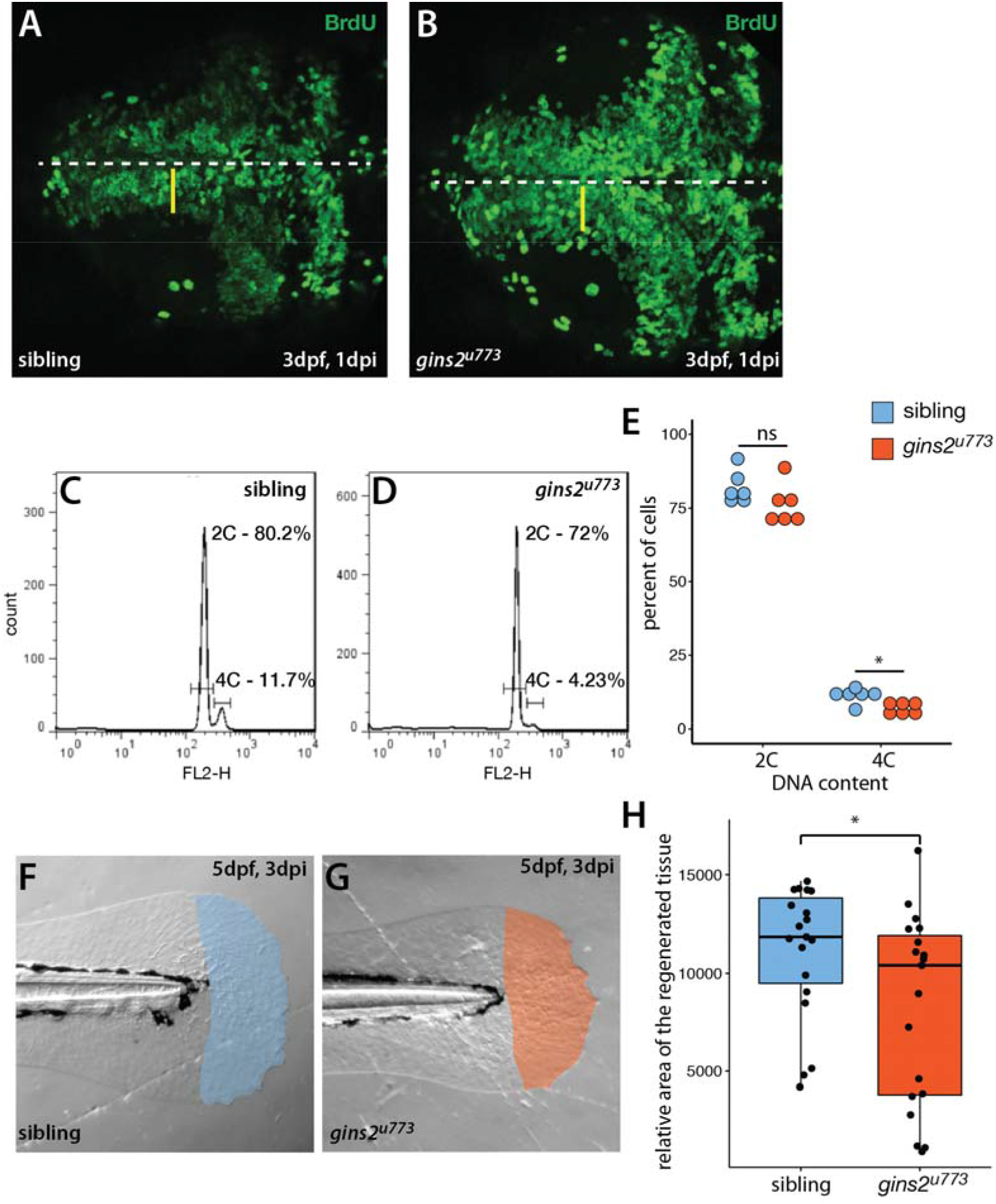
*gins2^u773/u773^* mutants show cell proliferation defects. A, B) The intensity of fluorescence of BrdU labeling in the OT of a 3 dpf wild type embryo shows a medio-lateral gradient. In contrast, mutant embryos showed a uniform distribution of the BrdU label. Dorsal views of the OT are shown, with anterior to the left. Midline is marked by the dashed line, yellow bars denote the proliferative area with a medial to lateral gradient of fluorescence in wildtype. C, D) Representative histograms of flow cytometry analysis of wildtype and mutant samples. E) DNA content analysis of propidium-iodide-labeled cells isolated from the head region showed no obvious change in the number of cells with 2C (p=0.15, n=6), and a slight, but significant decrease in the number of cells with 4C (p=0.01, n=6 independent samples). F, G) Regenerated area of the caudal fin in wildtype siblings and mutant embryos following amputation at 2 dpf. H) At 3 days post injury (dpi) the size of the regenerated fin fold was slightly but significantly smaller in mutant larvae than in controls (p=0.038, n=19).

To address if other processes requiring high levels cell proliferation are compromised in *gins2* mutants, we assessed fin regeneration in zebrafish larvae. We cut-off the distal tip of the larval caudal tail fin and measured the efficiency of regeneration, as defective regeneration would suggest that the proliferation of the cells required for fin regrowth is impaired. Three days after the original injury we observed a mild but significant decrease in the size of the *gins2^u773^* mutant regenerates (Figure 5F-H). Consequently cell division is likely compromised but not completely blocked during regeneration in *gins2^u773^* mutants.

### Structure of the GINS complex in the presence of Gins2^L52P^

To understand the structural consequences of the Gins2^L52P^ mutation, molecular dynamics simulations were carried out for the wildtype and mutant versions of homology models of the zebrafish Gins-complex (see Supplementary Figure 6). Surprisingly, the bH2 helix (naming convention in (Kamada et al., 2007)) of Gins2, which carries the L52P mutation, was not significantly destabilized.

Specifically we found that although the average helicity of residues 47-55 decreased from 69.1% to 49.1% in our simulations, the backbone trace of this segment in the mutant complex showed below average deviation (0.54 times the average backbone shift) from that of the wildtype structure. To further explore how the L52P switch altered the Gins complex, we examined the molecular neighborhood around Leu52. This amino acid is situated in a hydrophobic pocket lined by Ile17, Phe21, Ile26, Leu33, Phe36, Val44, Leu48 and Cys58 of Gins2 and Leu67 of the aH3 helix of Gins4. When Leu52 of Gins2 is switched to the smaller and less hydrophobic Pro, Leu67 of Gins4 moves closer, pulling several polar amino acids with it (Figure 6). Therefore, the structural change elicited by the Gins2^L52P^ mutation is predicted to strengthen the interaction between these two helices in the Gins2 and Gins4 proteins since the number of H-bonds within the region increases from a typical value of 2 to 4 as the result of the mutation (see Supplementary Figure 6).

In addition to stabilizing the interaction between these two helicies, the Gins2 L52P change induces a slight tilt of the aH3 helix in Gins4 that propagates to other domains of Gins4. One of the most significantly affected regions was the Gins4 aH2 helix, at the outermost surface of the complex, with a backbone shift 1.45 times that of the overall average. Structural studies have previously shown that the bH2 helix of Gins2, the tip of the aH2 and aH3 helices of Gins4 and the N-terminal segment of Gins4 form an organizational hot-spot in the active replisome complex, establishing direct contacts with both Cdc45 and Ctf14. It is likely, therefore, that even modest rearrangements in this region, such as those induced by the Gins2 L52P mutation, could impede replisome function. The nature of the interaction between Gins4 N-terminus and Ctf4 has been studied in detail in case of the yeast proteins. The N-terminal region of yeast Gins4, folded from its unstructured form into a helix by the interaction, fits into the binding-groove of the Ctf4, just as a very similar motif of Polα does establishing connection between CGM helicase and Polα (Simon AC et al., 2014, Villa et al, 2016). The yeast sequence contains a CIP-box motif within this highly flexible segment that shows capacity to form an α-helix (Villa et al., 2016), which is absent both in zebrafish and human gins4 sequences. However, in our simulations, the N-terminal 1-15 segment of zebrafish Gins4 also showed a propensity toward helix formation (forming a helix – even in the absence of Ctf4 – in 15% of the structures of the equilibrium ensemble), indicating a conserved mode of interaction between Gins4 and Ctf4.

**Figure 6.**
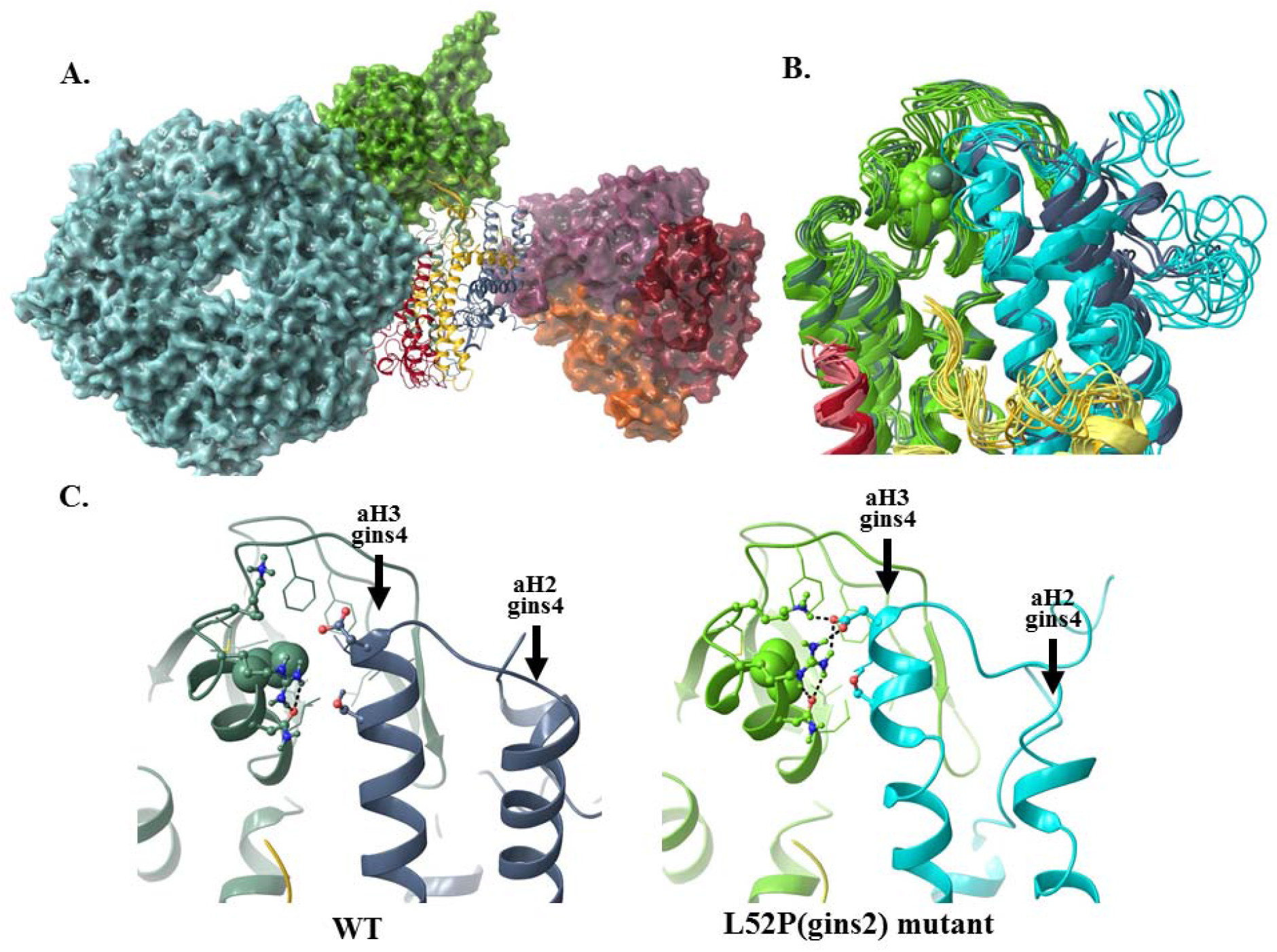
The structural consequences of the Gins2^L52P^ mutation in the zebrafish GINS complex. A) *In silico* generated model of the MCM-Cdc45-GINS-Ctf4 complex built using structures 3jc5 (Yuan et al., 2016), 4c8h (Simon et al., 2014) and the homology model of the zebrafish GINS complex created in this study. The MCM hexamer (cyan), Cdc45 (green) and the Ctf14 homotrimer (purple, red and orange) are shown as surfaces, while GINS as a ribbon diagram (Gins1: yellow, Gins2: green, Gins3: red, Gins4 blue). B) Superposition of the most populated cluster mid-structures (accounting for >90% of all structures of the equilibrium trajectory). Coloring of the Gins monomers is the same as before, darker shades are used for the wildtype, the lighter for the mutant structures (residue 52 of Gins2 shown in space-filling representation). C) Detailed view of the site of the mutation, with a typical H-bonding motif also illustrated.

## Discussion

### Loss of GINS function results in retinal and tectal apoptosis

Our work identifies the Gins2^L52P^ substitution caused by the *gins2^u773^* allele as a likely functional null mutation. Through phenotypic description of *gins2* and *gins1* loss-of-function mutations in zebrafish, we show that impairment of the GINS complex results in tissue-specific elevation of programmed cell death. Most likely due to defects in DNA replication, cells of the retinal and tectal proliferative zones undergo apoptosis whereas many other cells appear to be able to divide and differentiate normally.

Mutants for CMG components that associate with the GINS complex show similar, localized apoptosis phenotypes (Ryu et al., 2005) but there do appear to be some differences in phenotype upon loss of function of different members of the CMG complex. For instance, in the *mcm5* mutant, cells seem to be able to progress through S-phase to G2 whereas in the *gins2* mutant, our results suggest that cells appear to arrest early in S-phase, prior to completion of DNA replication. It is surprising that cells lacking *mcm5* behave differently to those with disrupted *gins2* function, as both encode components of the canonical CMG complex and the apoptotic phenotype in both mutants is observed in the same proliferative domains. However, in the case of the zygotic *mcm5* impairment, a high level of maternally provided Mcm5 protein could account for the late effects of the mutations (Ryu et al., 2005). Given that zebrafish embryos lack of the maternal Gins2 proteins, this cannot be the case in *gins2^u773/u773^* embryos. Therefore we propose that at early developmental stages and in some cell types, other factors substitute for GINS function (see below).

### GINS is absent during pre-somitogenesis phases of development

The lack of maternal component for Gins proteins is unexpected and surprising, as it suggests that until somitogenesis stages, when the zygotic expression of *gins* genes commences, DNA replication during cell divisions may happen independently of GINS function. Previous proteomic studies have also suggested the simultaneous presence of maternally provided Mcm proteins and absence of GINS-complex members in zebrafish oocytes (Ge et al., 2017). A potential explanation for the lack of survival in the absence of the GINS complex in the early embryo could be the peculiar nature of the cell cycle before the midblastula and early gastrula transitions (MBT and EGT, respectively) (Langley et al., 2014). During these stages of development, especially before MBT, cell cycles are extremely fast due to the lack of G phases (both G1 and G2 before MBT and G1 before EGT). The expression of *gins* genes can be observed only during somitogenesis (Thisse and Thisse, 2004; Thisse and Thisse, 2005) (Figure 3, Supplementary Figure 4) and in the absence of maternally deposited Gins proteins (Figures 3J, 4F), then it is possible that only after this stage is when “canonical” replication starts in the embryo. Prior to this, the MCM-complex may be able to perform its helicase function in the absence of GINS. Human dermal fibroblasts lacking GINS2 are also able to undergo mitosis, albeit at a slower rate, providing evidence that GINS-independent proliferation can also occur in mammalian cells (Barkley et al., 2009), and recent results suggest that other proteins, such as Fancm could substitute for GINS function under specific circumstances (Huang et al., 2019). It will be important to elucidate the identity of factors that can perform GINS-like function in young zebrafish embryos as they might be important, yet unrecognized, conditional components of the vertebrate replisome.

### The GINS-complex as possible target of cancer-therapeutics

The *gins2^u737^* missense allele provides new structural insights into the function of the eukaryotic replisome. In the CMG complex, Gins2 interacts directly with Gins1, Cdc45 and Mcm5 (Sun et al., 2015) and – through Gins4 – can also affect the conformation of Ctf4. Ctf4, in turn, provides the cross-link between the CMG helicase and the Polα/primase complex (Simon et al., 2014; Villa et al., 2016). Our *in silico* structural predictions indicate that the L52P change in Gins2 causes a strong rearrangement at the outermost surface of Gins4 near its N–terminus. This rearrangement may result both in altering both the GINS-Cdc45 interaction and also disrupting the interconnection between CMG and Polα.

As we find targeting Gins2 function predominantly compromises proliferation and viability of fast proliferating cells, we propose that the interaction between Gins2 and Gins4 could be exploited as a therapeutic target to inhibit such cells in malignancies. The components of the GINS-complex have previously been identified as important biomarkers (Kanzaki et al., 2016; Tauchi et al., 2016; Yamane et al., 2016) and in accordance with their potentially important role as a therapeutic target, several studies show that in the absence of GINS function, the proliferative and invasive capacity of cancer cells decreases (Liang et al., 2016; Yamane et al., 2016; Zhang et al., 2015; Zhang et al., 2013). The inhibition of the replisome in cancer therapy through the targeting of the Mcm2-7 helicase has also been suggested (Simon and Schwacha, 2014), and we propose that the impairment of other CMG components is a logical extension of this idea.

The fact that the impairment of the GINS-complex does not appear to affect slow-cycling cells and that early embryonic cells do not need GINS to successfully proliferate suggests that there might be other, yet to be described, non-canonical forms of the replisome. If the existence of such GINS-free replication machineries in multiple cell types can be proven, it would also enhance the therapeutic potential of the GINS complex, as its disruption might have consequences restricted to fast proliferating cell populations.

### Possible pleiotropic functions of GINS-complex components

Although the role of *GINS* genes in the replisome is clearly established, there is tantalizing evidence that they could perform other functions as well during development. For example, GINS2 has been shown to have a proliferation-independent role in the mitotic segregation of chromosomes (Huang et al., 2005) and a possible role for *GINS3* in cardiac physiology has been pinpointed by several genome-wide association studies (GWAS) (Arking et al., 2014; Milan et al., 2009; Verweij et al., 2016). In our analyses, we observed disorganization of the OT and the retina, though it remains to be determined if these are due to other, pleiotropic roles of Gins2 or a secondary consequence of proliferation and apoptosis defects in these regions.

Further studies are necessary to establish the spectrum of pleiotropic effects upon disruption of GINS components and we anticipate that genetic models in zebrafish will provide important insights to both replisome dependent and independent functions of these conserved proteins.

## Methods

### Fish husbandry and mutagenesis

Wildtype and mutant fish lines were maintained in the animal facilities of University College London and ELTE Eötvös Loránd University according to standard protocols (Aleström et al., 2019; Westerfield, 2000). All protocols used in this study were approved by the UK Home Office and the Hungarian National Food Chain Safety Office (Permit Number: XIV-I-001/515-4/2012). Mutagenesis was performed in wildtype male AB/TL strain fish by ENU treatment as described previously (van Eeden et al., 1999; Young et al., 2019). Fish stocks were maintained by outcrossing heterozygous carriers to wildtype *ekwill (ekw)* fish.

### Genetic mapping, morpholino injections and creation of a novel *gins1* loss-of-function allele

The *gins2^u773^* allele was mapped by bulk segregant analysis with SSLPs to L18 (Talbot and Schier, 1998). Markers z7256 and z10008 flanked a ~12.6 Mb interval containing 183 genes. Next generation sequencing, combined with a slightly modified version (Turner et al., 2019) of the Variant Discovery Mapping (VDM) CloudMap pipeline (Minevich et al., 2012) pipeline confirmed and refined this position. Instead of plotting individual allele frequencies, the kernel density of homozygous/heterozygous SNPs was plotted along each chromosome. From the list of variants in this mapping region, background SNPs identified through the sequencing of the *ekwill* strain plus a list compiled from previously published data (LaFave et al., 2014; Obholzer et al., 2012) was subtracted leaving a total of 13 candidate protein coding variants. After eliminating genes that are not expressed during early development and/or do not have mutations in conserved regions, we were able to narrow down our search to *gins2.*

FASTQ files are available at the SRA, …..

Primers used to amplify the genomic region within *gins2* relevant to the *u773* allele with Sanger sequencing: gins2-ex2-F – 5’-GAGATTTAGGCCCGTTCAACCC-3’ and gins2-ex3-R – 5’-TCTCCTGTTCACGGATGTCTTCT-3’.

The splice site specific synthetic MO that was used to knock-down *gins2* function was purchased from Gene-Tools (Philomath, OR, USA). The sequence of the *gins2MO* was: 5’-GGGGTGAGTCAATTTATAATCTAC-3’. In total 1 nl of 1 mM *gins2MO* was injected per embryo at the one cell stage of development.

To induce a mutation in the *gins1* gene we used the CRISPR/Cas9 system as described (Gagnon et al., 2014). The targeted site within the 4^th^ exon of the gene (with the PAM sequence in bold): GGGCACGGATTCGCAAGAGGCGG. The primers used to amplify the respective genomic region of the *gins1* gene were: gins1-ex3-F – 5’-GTAGGAACGAGGCAAAGACAGAGG-3’ and gins 1-in4-R – 5’-TGGGCTTGATCTTCTGTGTAGC-3’.

### *In situ* hybridization, immunohistochemistry, TUNEL staining and BrdU analysis

A detailed protocol for *in situ* hybridization has been described before (Bellipanni et al., 2006). *gins2* cDNA was cloned using primers gins2-F – 5’-CTCCTTGACGTCAGAGACACAT-3’ and gins2-R – 5’-GGAGAGGAATGGCTGAAGTACC-3’ into pCR-Blunt II-TOPO vector (Invitrogen) following the manufacturer’s protocol, cut with KpnI and transcribed using SP6 polymerase.

TUNEL labeling to detect apoptotic cells was performed using ApopTag kits (EMD Millipore), following manufacturer’s instructions and a previously described protocol (Cerveny et al., 2010). Apoptotic cells were detected in situ with a Peroxidase-coupled Digoxygenin antibody or with a Rhodamine-coupled Digoxygenin antibody.

Proliferating cells were detected using a BrdU pulse-chase protocol. At 2 dpf embryos were anesthetized, and embedded into 0.8% low-melting point agarose in E3. Approximately 2 nl of 2.5ng/μl BrdU was injected into each heart. Embryos were recovered from the agarose, and moved into fresh E3 medium. At 3 dpf, embryos were fixed in 4% paraformaldehyde and stained with an anti-BrdU antibody as described (Cerveny et al., 2010).

TUNEL or *in situ* stained embryos were embedded in a JB-4 resin according to the manufacturer’s protocol. Sectioning was done with a Leica RM2025 microtome and 10 μm sections were collected.

### DNA content analysis

Heads of 10 mutant and 10 wildtype 2 dpf anaesthetised embryos were dissected using a sharp tungsten needles. Single cell suspensions were generated after room temperature incubation for 30 min in 20 units/ml papain (Sigma-Aldrich, P4762) in L15 tissue culture media (Sigma-Aldrich, L5520). Cells were washed in PBS and fixed in ice-cold 70% (v/v) ethanol and stored overnight at −20°C. Cell pellets were resuspended in 0.5 ml hypotonic fluorochrome solution (4 mM citrate buffer, pH 6.5, containing 50 μg/ml propidium iodide (Sigma-Aldrich), 200 μg/ml RNase A (cat. no. 12091-021), 0.1% Triton X-100) and kept in the dark at 4°C until analysis. Data acquisition was performed by using a FACSCalibur flow cytometer (Becton Dickinson). Debris were excluded by the FSC-SSC dot-plot and the percentage of diploid (2C) and tetraploid (4C) cells were determined by the FlowJo software (FLOWJO, LLC).

### Western blot analysis

For Western blot analysis 10 embryos were lysed in 2x Laemmli reducing sample buffer (Bio-Rad 1610737), sonicated for 3 min and separated by SDS-PAGE and transferred to nitrocellulose membranes. Membranes were immunoblotted using antibodies against Psf1 (Origene, TA339351) Psf2 (Novus Biologicals, NBP1-58209) and γ-tubulin (Sigma, T6557). Bindings were revealed by a HRP-conjugated anti-mouse/rabbit IgG antibodies (Dako) and ECL (Advansta, WesternBright) detection.

### Cell transplantations

Cell transplantations were performed similarly to previous work (Cerveny et al., 2010). Donor embryos were generated by injecting one-cell stage embryos with GFP mRNA (approximately 25-30 pg per embryo) followed by 1nl of 1 mg/ml *gins2MO* or no additional material. Approximately 40 GFP+ cells were transplanted from the animal pole of mid-blastula stage embryos into an equivalent area in similarly staged host embryos. Embryos were immobilized in 4% methylcellulose in embryo medium and viewed at 40X magnification with a compound microscope (Nikon E1000). Cells were moved by suction using an oil-filled manual injector (Sutter Instrument Company). Donor and host embryos were incubated overnight in E3+1% Penicillin+Streptomycin (Sigma) and then incubated in E3+PTU until fixation with 4% PFA at 3 dpf. After fixation, embryos were washed with PBST, dehydrated in a graded MeOH series, and stored at −20°C for at least 2 days until TUNEL staining. n≥16 transplants of morpholino-injected and control donors.

### Cartilage staining and measurements

Cartilage staining was performed as described (Westerfield, 2000). Briefly, larvae fixed in 4% paraformaldehyde were dehydrated in an ethanol series, and stained in an alcian blue solution overnight, at room temperature. Pictures taken from the ventral side of specimens were analyzed with ImageJ and statistical analysis was done with RStudio.

### Tail regeneration assay

Caudal tail fins, posterior to the notochord of 2 dpf anaesthetised mutant and sibling embryos were amputated using a sharp tungsten needle. Pictures were taken right after the intervention and the individual larvae were placed in 24 well plates, where they remained until 5 dpf. Pictures were taken of the regenerated tail fin tissue and using the 2 dpf pictures as references, the approximate size of the regenerate was calculated using ImageJ. Statistical analysis was performed using RStudio.

### Molecular Modeling

A homology model of the zebrafish Gins complex was built using structures of the human complex (2e9x (Kamada et al., 2007), 2q9q (Chang et al., 2007)) utilizing the sequence identities of 83%, 81%, 67% and 69% in case of Gins1, Gins2, Gins3 and Gins4, respectively (sequences were taken from F1RAD9, Q4VBJ6 and F1QXM4 entries of Uniprot for of Gins1, Gins2, Gins3 and ZDB-GENE-040801-59 for Gins4). Homology modeling, building missing segments (1-18 of Gins4) and initial minimizations were carried out using the Schrödinger Suite (Schrödinger Release 2016-4: Maestro, BatchMin, QSite, Schrödinger, LLC, New York, NY, 2016.). Wildtype and mutant structures were then subjected to extensive molecular dynamics simulation, using GROMACS 5.0 (Pronk et al., 2013) and the AMBER99SB-ILDNP forcefield (Aliev et al., 2014). Systems were solvated by TIP3P water molecules in dodecahedral boxes with 10Å buffer. The total charge of the system was neutralized and physiological salt concentration set using Na^+^ and Cl^-^ ions. Energy minimization of starting structures was followed by sequential relaxation of constraints on protein atoms in three steps (all of 100ps). Trajectories of 1000 ns NPT simulations at 325 K and 1 bar were recorded for further analysis (collecting snapshots at every 4ps). Averaging and clustering of conformations (Daura et al., 1999) was carried out based on the main-chain conformation of snapshots from the last 200ns of the simulations (which is referred to as “equilibrium trajectory”) using a cutoff of 1.5 Å. The C-terminal segment of Gins1 (residues 145-196, its entire B-domain) was shown to be unstructured in the uncomplexed form (Chang et al., 2007), thus it was not included in the simulations, but was attached to the final structures (modeled using the yeast gins-CGM complex (3jc5 (Yuan et al., 2016), 42% identity for the gins1 B-domain) followed by a thorough energy minimization.

### Multiple sequence alignments, data analysis and visualization

Statistical analysis, multiple sequence alignments and visualization was performed in R (R Core Team, 2018) using the *msa* and *ggplot2* packages (Bodenhofer et al., 2015; Wickham, 2016).

## Acknowledgments

We thank Carole Wilson and her team for fish care. This work was funded by grants from the Biotechnology and Biological Sciences Research Council [BB/H008462/1 to S.W.W.], Medical Research Council [MR/L003775/1 to S.W.W. and Gaia Gestri], Wellcome Trust [095722/Z/11/Z and 207483/Z/17/Z to R.J.P, and 089227/Z09/Z and 104682/Z/14/Z to S.W.W.], OTKA [K116305 to D.K.M.] and VEKOP-2.3.3-15-2016-00007. M.V. was also supported by the ÚNKP-17-4 New National Excellence Program and the ELTE Institutional Excellence Program (1783-3/2018/FEKUTSRAT) both sponsored by the Hungarian Ministry of Human Capacities.

## Author Contributions

Conceptualization: Máté Varga, Stephen W. Wilson.

Data curation: Richard J. Poole.

Funding acquisition: Stephen W. Wilson, Dóra K. Menyhárd, Máté Varga.

Investigation: Máté Varga, Kitti Csályi, Dóra K. Menyhárd, Richard J. Poole, Kara L. Cerveny, Dorottya Kövesdi, Balázs Barátki, Hannah Rouse, István Bertyák, Zsuzsa Vad, Rodrigo Young, Thomas Hawkins, Heather Stickney, Florencia Cavodeassi, Quenten Schwarz.

Methodology: Máté Varga, Kara L. Cerveny, Richard J. Poole, Dóra K. Menyhárd, Dorottya Kövesdi, Rodrigo Young.

Project administration: Máté Varga, Stephen W. Wilson.

Supervision: Máté Varga, Stephen W. Wilson.

Writing – original draft: Máté Varga, Dóra K. Menyhárd, Kara L. Cerveny, Richard J. Poole, Stephen W. Wilson.

