## Supplemental Figure for "The GINS complex is required for the survival of rapidly proliferating retinal and tectal progenitor cells during zebrafish development"

### The GINS complex is required for the proliferation of neural progenitors during zebrafish development

#### Supplementary Figures and Tables

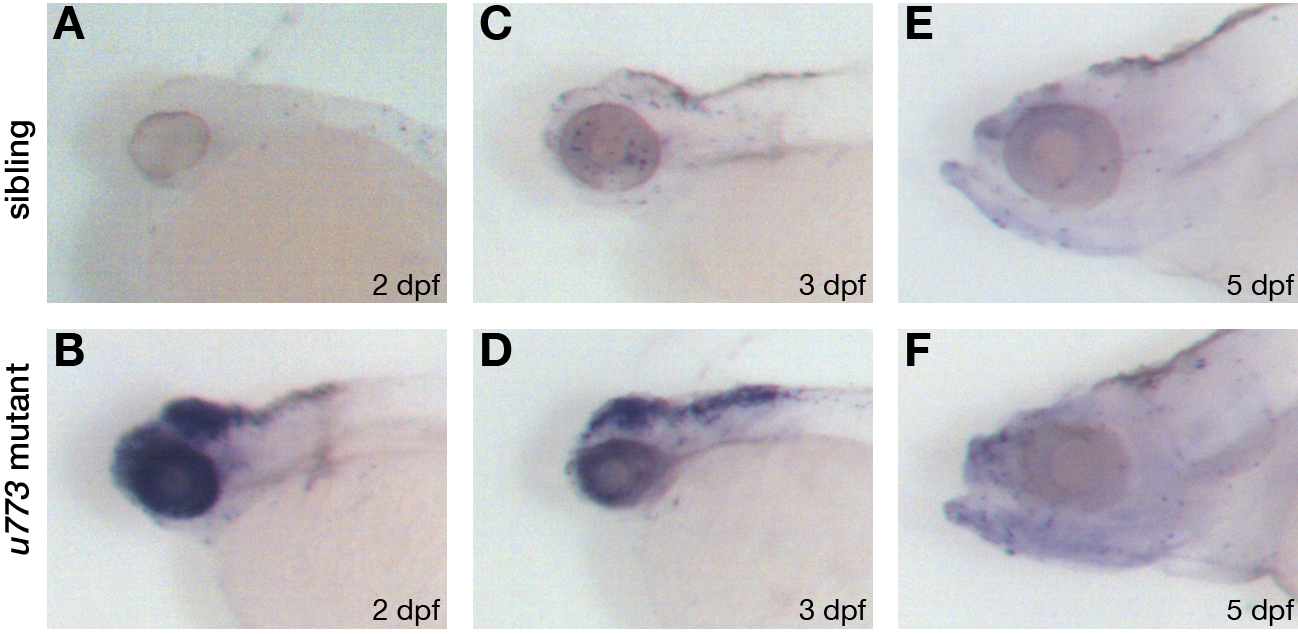

Supplementary Figure 1. Transient apoptosis can be observed in *u773* mutant embryos**.**

A-F) Lateral views of TUNEL-stained wildtype (A,C,E) and *u773* mutant (B,D,F) embryos at stages indicated.

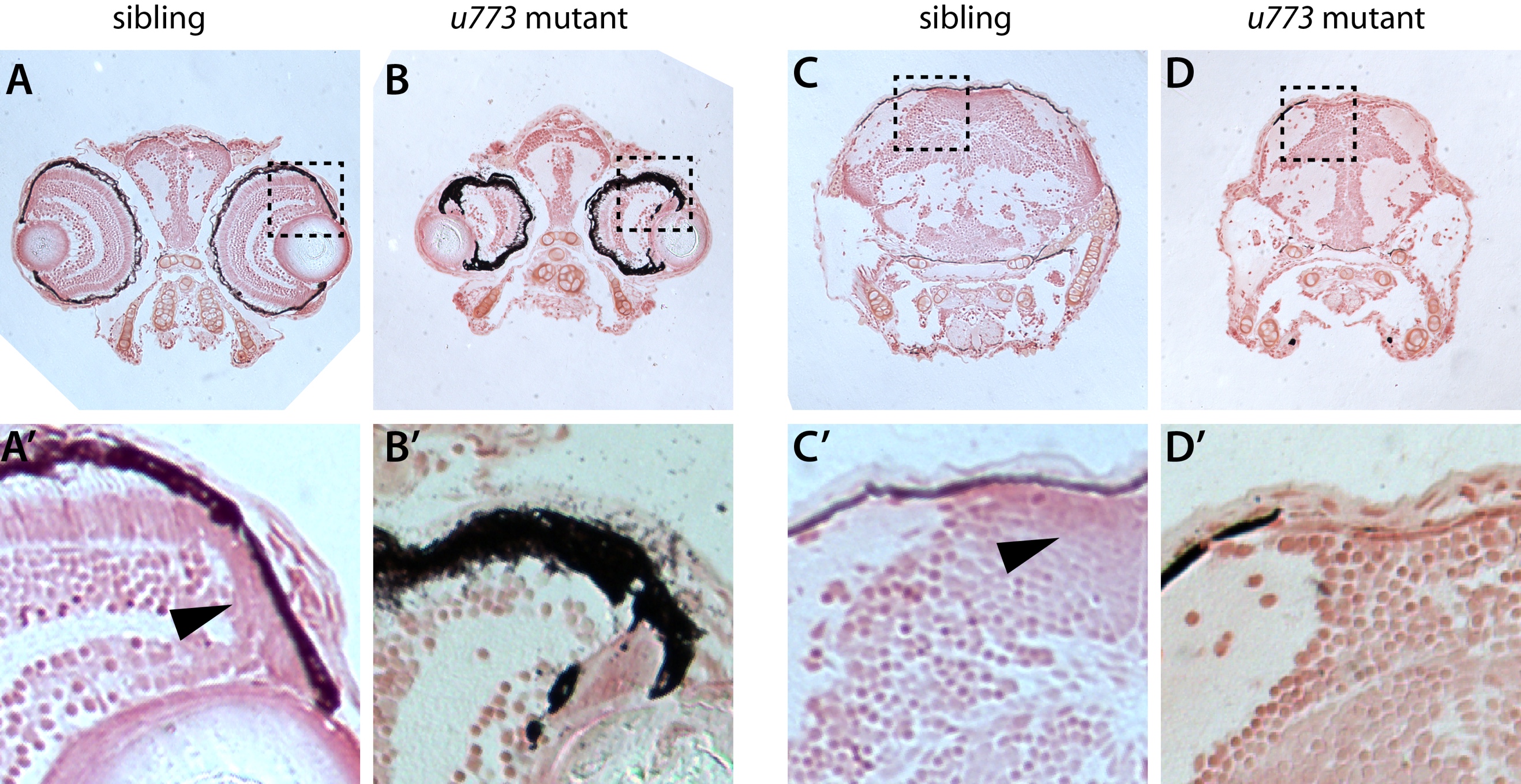

Supplementary Figure 2. Histological analysis of 5 dpf *u773* mutant larvae.

A-B’) Transverse sections of wildtype and mutant zebrafish larvae through the eyes. A’ and B’ show magnified views of the boxes in A and B. Note that the progenitor zone in the eye, denoted by the arrowhead in the wildtype larvae (A’) is missing in the *u773* mutants (B’).

C-D’) Transverse sections of wildtype and mutant zebrafish larvae through the OT areas. C’ and D’ show magnified views of the boxes in C and D. Note that the progenitor zone in the dorsomedial region of the tectum, denoted by the arrowhead in the wildtype larvae (C’) is missing in the *u773* mutants (D’).

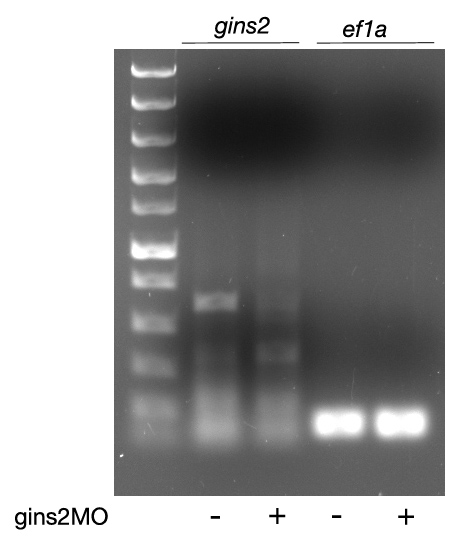

Supplementary Figure 3. Injection of gins2MO reduces *gins2* mRNA levels.

RT-PCR on 2 dpf *gins2*MO injected samples suggests that mis-spliced *gins2* mRNA undergoes non-sense mediated decay (NMD).

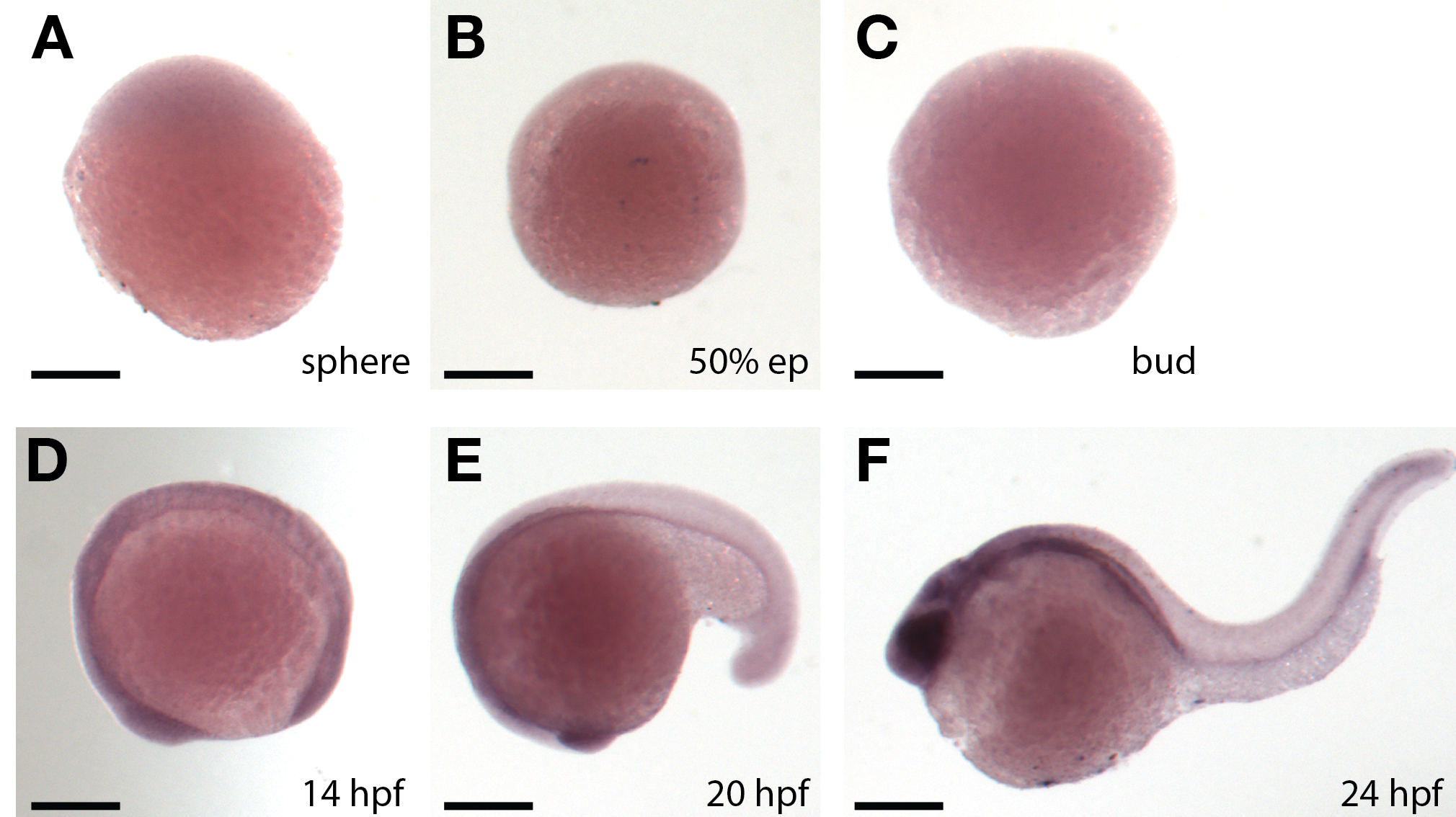

Supplementary Figure 4. The expression of *gins1* during zebrafish development.

A-F) Lateral views of the wildtype embryos assessed for *gins1* expression with whole mount *in situ* hybridization at the indicated stages. Scale bars: 150 μm.

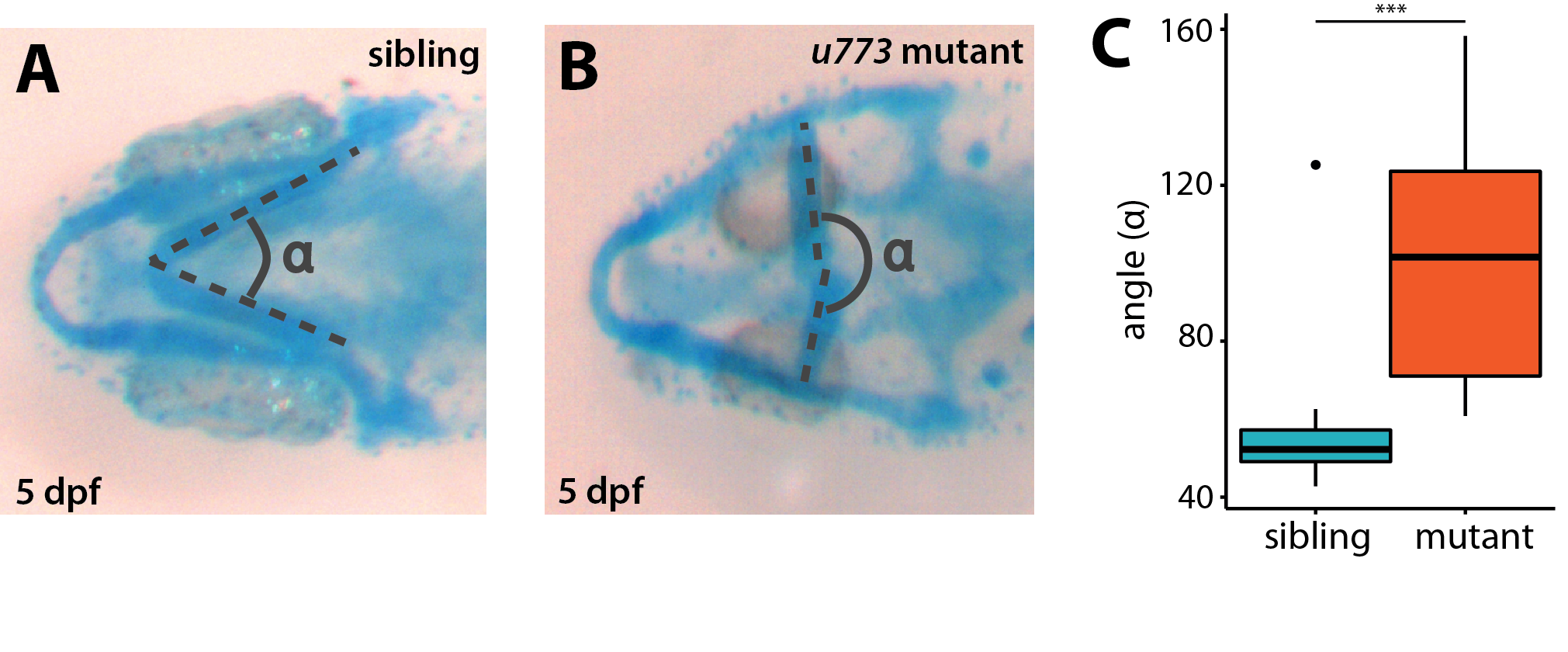

Supplementary Figure 5. Jaw cartilage development is severely impaired in *u773* mutants.

A,B) Alcian Blue staining of the jaw cartilage in 5 dpf wildtype (A) and *u773* mutant (B) larvae.

C) The angle of ceratohyals of 5 dpf mutant larvae stained is significantly different compared to wildtype (n=19) (p<0.001).

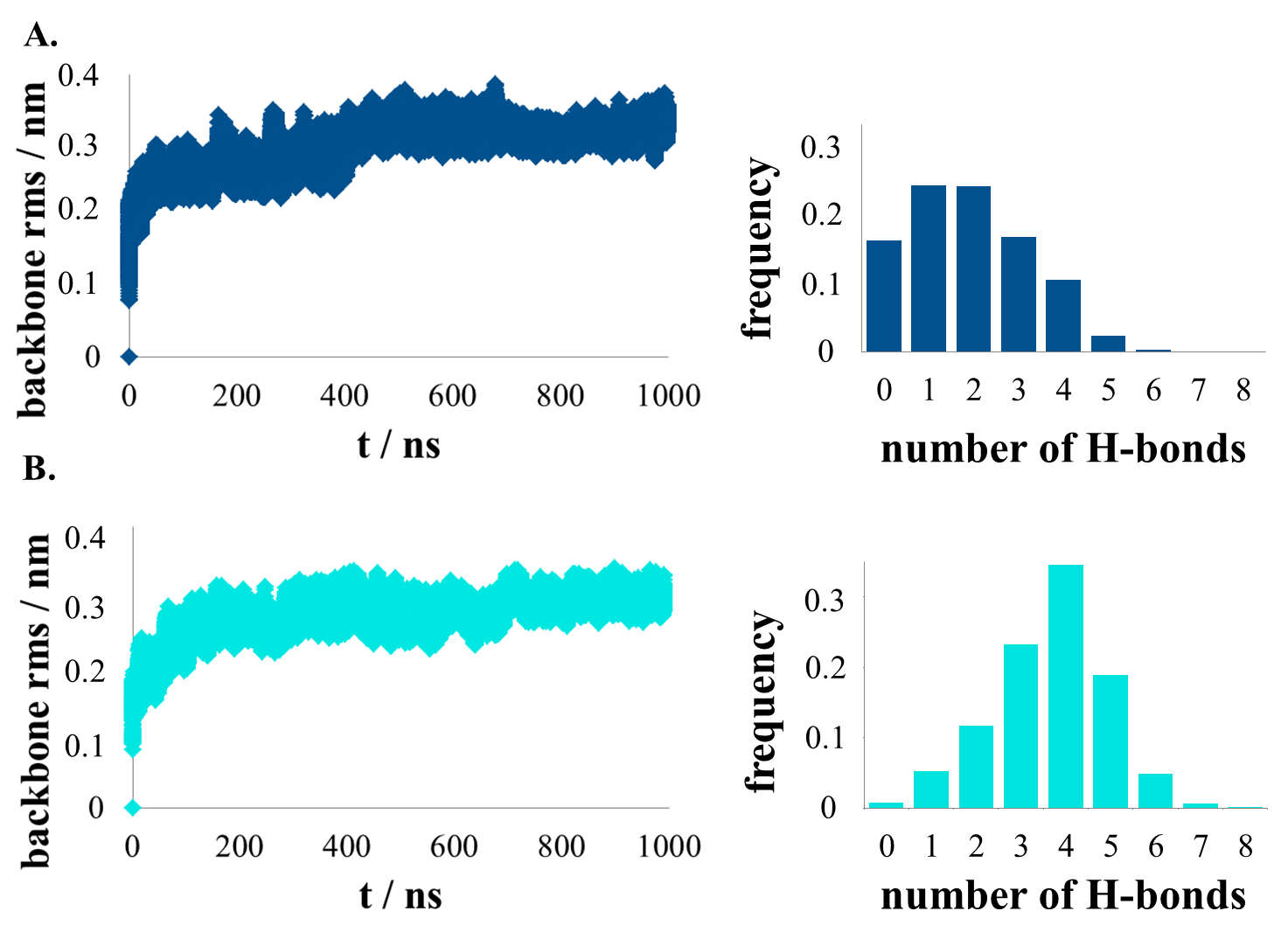

Supplementary Figure 6. The root-mean-square deviation of the protein backbone and the number of H-bonds formed between the neighboring regions of Gins2 (residues 48-58) and Gins4 (64-74) in case of the wildtype (A) and the Gins2^L52P^ (B) tetramer.

**Supplementary Table 1.** (Dóra)

#### Supplementary Materials and Methods

**Neutral Red staining**

Embedding and sectioning of 5 dpf larvae was performed as described in the general Materials and Methods section. Sections were stained in freshly prepared 0.001% Neutral Red (N4638, Sigma-Aldrich) solution for 5 min, washed in dH_2_O, dried and covered.

**Alcian Blue staining**

Larvae were fixed in 4% PFA, incubated in 50% ethanol for one day and transferred to 100% ethanol. After 24 hours, the staining solution was added (0.02% Alcian Blue (A5268, Sigma-Aldrich), 29.8% acetic acid, 70% ethanol)
